# Direct Energy Transfer from Allophycocyanin-Free Rod-Type CpcL-Phycobilisome to Photosystem I

**DOI:** 10.1101/2021.05.31.446472

**Authors:** Tomoyasu Noji, Mai Watanabe, Takehisa Dewa, Shigeru Itoh, Masahiko Ikeuchib

## Abstract

Phycobilisomes (PBSs) are photosynthetic antenna megacomplexes comprised of pigment-binding proteins (cores and rods) joined with linker proteins. A rod-type PBS that does not have a core is connected to photosystem I (PSI) by a pigment-free CpcL linker protein, which induces a red-shift of the absorption band of phycocyanobilin (PCB) in the rod (red-PCB). Herein, the isolated supercomplex of the rod-type PBS and the PSI tetramer from *Anabaena* sp. PCC 7120 were probed by picosecond laser spectroscopy at 77 K and by decay-associated spectral analysis to show that red-PCB mediates the fast (time constant = 90 ps) and efficient (efficiency = 95%) transfer of excitation energy from PCB in rod to chlorophyll *a* (Chl *a*) in PSI. According to the Förster energy transfer mechanism, this high efficiency corresponds to a 4-nm distance between red-PCB and Chl *a*, suggesting that β-84 PCB in rod acts as red-PCB.

**TOC GRAPHIC:** 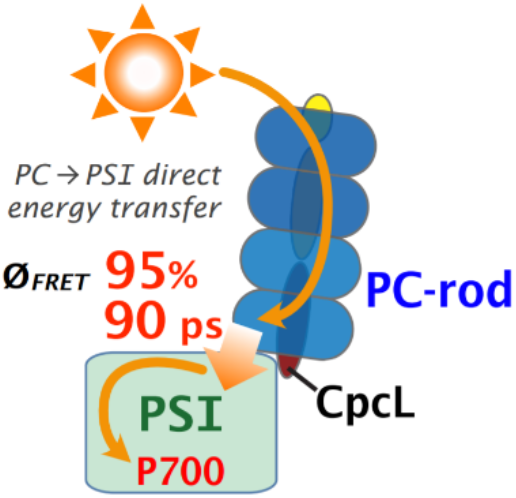

---

Oxygenic photosynthesis is enabled by the sequential operation of two types of photosystems (PSs), namely, PSI and PSII, which accumulate the energy of sunlight absorbed by pigments of the peripheral antenna and the reaction center (RC), and use it to promote the photoreactions of chlorophyll *a* (Chl *a*) in the central cores of the RCs of both PSs.^1^ In cyanobacteria, multiple types of phycobilisomes (PBSs) harvest the green/orange part of sunlight and transfer the excitation energy to PS-attached Chl *a*, which mainly absorbs blue and red light.^1^ Typical PBSs (rod-core-type PBSs) are large extra-membranous pigment-protein complexes comprised of central cores and peripheral rods. Basically, two or more phycocyanin (PC) hexamer disks extending from the central allophycocyanin (APC) core.^1^ Both PC and APC contain the same pigment, phycocyanobilin (PCB). The absorption/fluorescence peaks of APC-bound PCB are red-shifted compared to those of PC-bound PCB (650–670/660–680 nm vs. 550–630/640– 650 nm, respectively).^1–3^ Therefore, the fluorescence band of PCB on PC fully overlaps with the red-shifted absorption band of PCB on APC, while the red-shifted fluorescence band of APC-bound PCB overlaps with the absorption bands of Chl *a* in PSs, peaking at 670–690 nm. These extensive overlaps explain the efficient PC → APC → PS energy transfer. Rod-core-type PBSs are known to transfer excitation energy mainly to PSII, although transfer to PSI is also possible in some cases.^4–8^ Some PBSs have simpler structures without the central core APC, as exemplified by those of the cyanobacterium *Acaryochloris marina*, which uses Chl *d* both in PSI and II, instead of Chl *a*, and relies on the transfer of excitation energy to Chl *a* in PSII through the PC/APC hetero-hexamer formed at the rod base.^9, 10^ APC has therefore been considered essential for the energy transfer from both rod-core- and rod-type PBSs to Chls on PSI/PSII. ^1^, ^4–10^

In the PBS-CpcL-PSI_4_ supercomplex, which has been recently discovered and isolated from the cyanobacterium *Anabaena* sp. PCC 7120 (hereafter *Anabaena*),^2^ a rod-membrane linker protein (CpcL) connects the PC rod to the PSI tetramer (PSI_4_) in the absence of APC.^11^ Notably, *cpcL* genes are widely conserved in cyanobacterial genomes^2, 12^ and are preferentially expressed under conditions requiring larger PSI excitation, e.g., in cases of nitrogen starvation or under green light conditions.^2, 12–14^ The energy transfer from CpcL-PBS to PSI_4_ has been suggested by steady-state fluorescence spectroscopy,^2^ however, the efficiency and time constant of the energy transfer from CpcL-PBS to PSI_4_ in the supercomplex have not been determined yet.

Herein, we prepared four types of samples (Figure S1A), namely, (i) CpcL-PBS, (ii) PSI_4_, (iii) the PBS-CpcL-PSI_4_ supercomplex, and (iv) a CpcL-PBS/PSI_4_ mixture with a 1:1 (mol/mol) ratio of CpcL-PBS and PSI_4_ (as observed in the native PBS-CpcL-PSI_4_ supercomplex; page S10, and Figures S1B and S2 together with their detailed descriptions). Figure 1 shows the two-dimensional (emission wavelength/time after laser excitation) fluorescence images of the isolated CpcL-PBS complex excited at 628 nm (Figure 1A) and the PBS-CpcL-PSI_4_ supercomplex excited at 405 and 628 nm (Figures 1B and C, respectively) measured at 77 K. The laser pulses at 405 and 628 nm almost selectively excited Chl *a* and PCB, respectively.^3, 15^ The fluorescence emission spectrum of CpcL-PBS excited at 628 nm featured two bands at (i) 650 nm and (ii) 669 nm (Figure 1A), which corresponded to the fluorescence bands of (i) PC-bound PCB (F645) that absorbs at 633 nm (A633) and (ii) CpcL-modified PC-bound red-shifted PCB (F669) that absorbs at 659 nm (see pages S13–S14 as well as Figure S3A), respectively.

**Figure 1.**
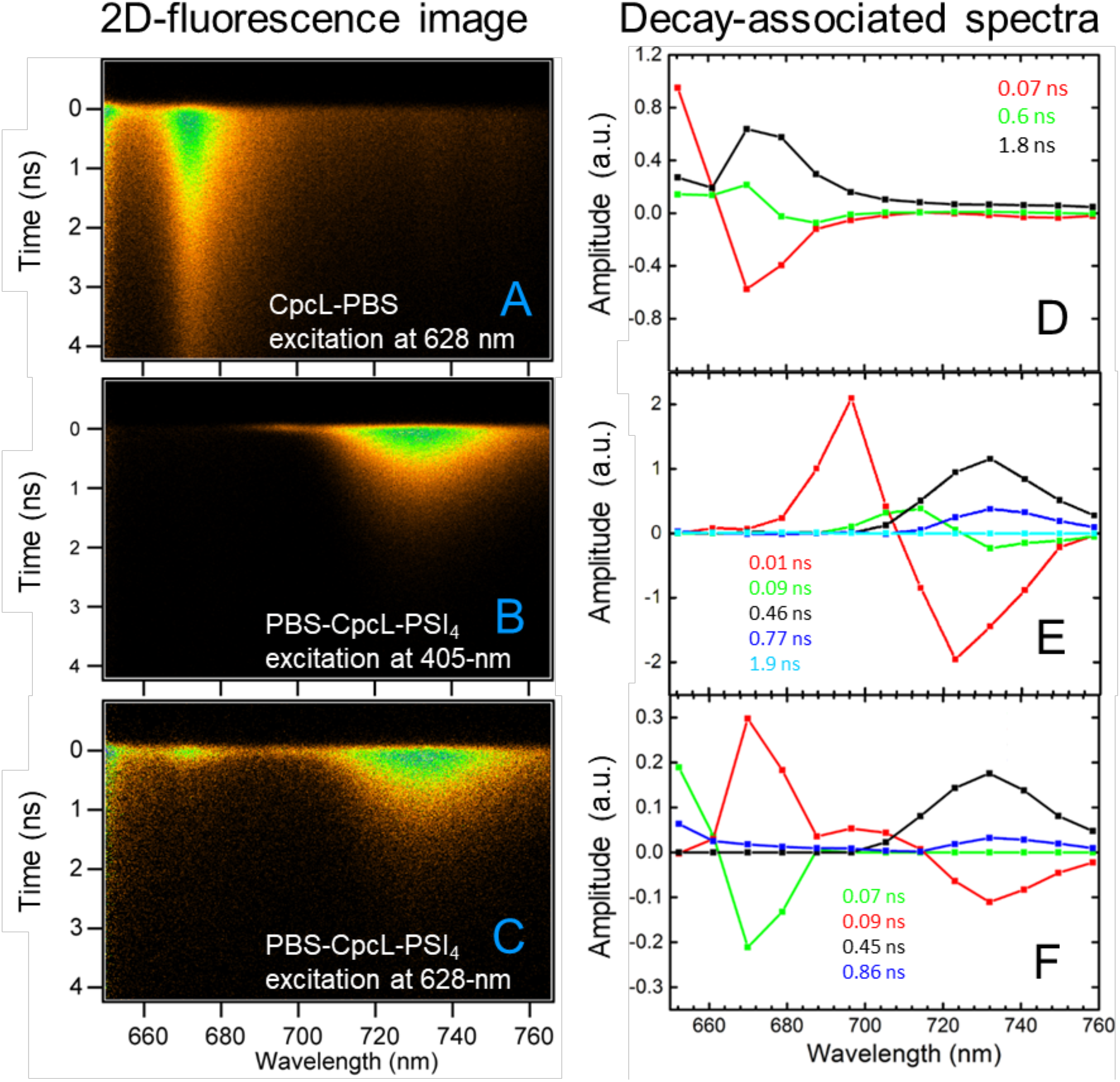
Two-dimensional fluorescence images measured at 77 K (A–C) and decay-associated spectra (DASs; with time constants) calculated from these images (D–F). (A, D) CpcL-PBS complex excited at 628 nm. (B, E) PBS-CpcL-PSI_4_ supercomplex excited at 405 nm. (C, F) PBS-CpcL-PSI_4_ supercomplex excited at 628 nm. For (A) and (C), fluorescence emission at wavelengths below 650 nm was eliminated using a 650-nm long-pass filter. The excitation laser intensities were set to 1 *μ*W.

For the PBS-CpcL-PSI_4_ supercomplex, irradiation with a 405-nm laser, which mainly excites Chl *a* in PSI_4_, induced only a broad 729-nm (F729) band (Figure 1B) that was absent in Figure 1A and could be assigned to the typical red-Chl *a* component on PSI_4_ (Figure S4). The excitation of the PBS-CpcL-PSI_4_ supercomplex at 628 nm induced strong F645 and F729 bands (Figure 1C) as well as a significantly weaker (compared to that in Figure 1A) F669 band. The F669 band in Figure 1C featured a short vertical tail, which indicated fast decay and, thus, a significantly shorter lifetime than that observed for CpcL-PBS alone. This result indicated efficient energy transfer from PCB (F645) to red-Chl *a* (F729) in PSI_4_ via red-PCB (F669).

The time course of fluorescence decay at each emission wavelength (Figure 1D–F and S5) was calculated from the images shown in Figures 1A–C. Compared to that for CpcL-PBS (1.8 ns) or the CpcL-PBS/PSI_4_ mixture (1.72 ns; Figures 1D and S5–6), the decay of F669 for the PBS-CpcL-PSI_4_ supercomplex was much faster, featuring an apparent time constant of 0.09 ns. The energy transfer from CpcL-PBS to PSI_4_, which was absent in the CpcL-PBS/PSI_4_ mixture, seemed to be very efficient in the PBS-CpcL-PSI_4_ supercomplex (see Figures 2 and S5–S7, as well as pages S17–S18).

**Figure 2.**
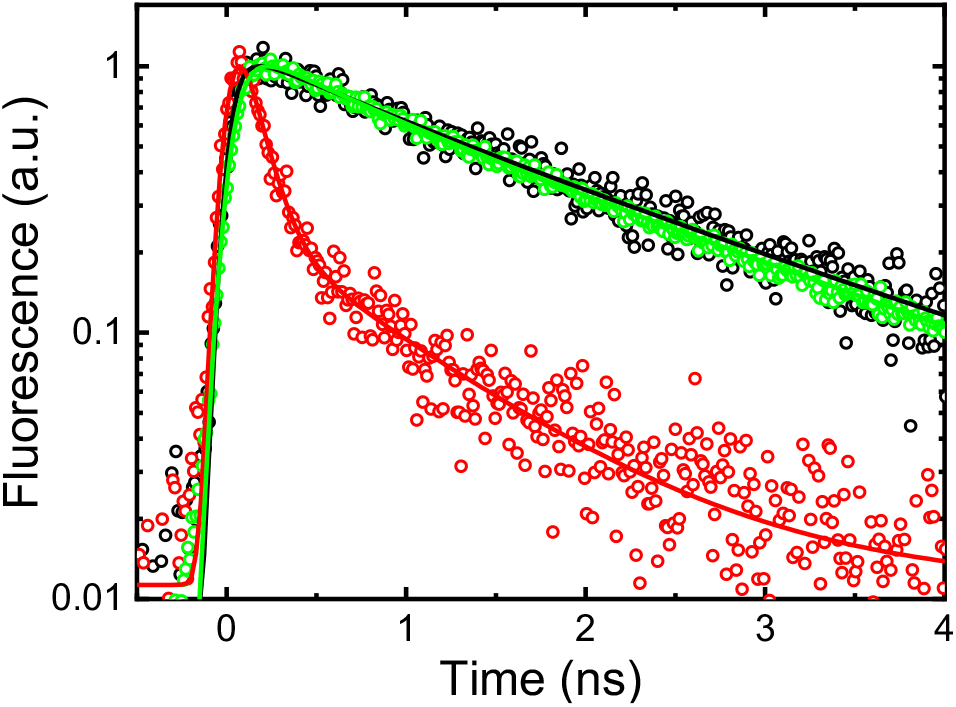
Time courses of fluorescence decay measured at 669 nm for CpcL-PBS (green circles), the PBS-CpcL-PSI_4_ supercomplex (red circles), and the CpcL-PBS/PSI_4_ mixture (black circles) after laser excitation at 628 nm. Solid lines represent best-fit curves normalized at peak intensities.

This result confirmed the detection of high F729 intensity for the steady-state fluorescence of the PBS-CpcL-PSI_4_ supercomplex excited at 550–650 nm (Figure S3B) and indicated an increase in the effective absorption cross-section of PSI_4_ in the supercomplex to include the absorption bands of PCB and red-PCB contained in CpcL-PBS.

Figures 1D–F show the decay-associated spectra (DASs) calculated by the global fitting (Figures S5) of fluorescence decay at various wavelengths, estimated from the images in Figures 1A–C. The negative and positive bands of each DAS curve represent the rise and decay of fluorescence intensity that feature the same time constant.

The fastest (time constant (*t*_c_) = 0.07 ns) decay curve that was calculated for the CpcL-PBS complex excited at 628 nm (Figure 1D) had a large positive peak at 645 nm (F645) and a negative peak at 669 nm (F669). This suggests a fast energy transfer from F645 (PCB) to F669 (red-PCB) in the CpcL-PBS complex. The positive bands at 670–680 nm in the slower (*t*_c_ = 0.6 and 1.8 ns) decay curves in Figure 1D suggest energy dissipation from F669.

For the PBS-CpcL-PSI_4_ supercomplex irradiated by a 405-nm laser (which mainly excites Chl *a* and very weakly excites PCB), a large positive F696 peak and a large negative F729 peak in the 0.01-ns decay curve suggest a fast 0.01-ns-scale energy transfer from F696 (Chl *a*) to F729 (red-Chl *a*) (Figure 1E). The 0.45-ns decay showed only a positive peak at 729 nm, indicative of energy dissipation from F729 in PSI_4_. This result was almost identical to that reported for the isolated *Thermosynechococcus elongatus* PSI trimer containing no PBS.^16^ Notably, all three DASs resembled those observed for the PSI_4_ complex alone, showing almost negligible contributions of the CpcL-PBS moiety in the shorter wavelength range (Figure S6). Therefore, the DAS curves in Figure 1E represent energy transfer from bulk antenna Chl *a* to red-Chl *a* inside PSI and the following dissipation of energy from red-Chl *a*.

The fastest decay curve of the PBS-CpcL-PSI_4_ supercomplex, excited at 628 nm (Figure 1F), showed a time constant of 0.07 ns (Figure 1C). The time constant is comparable to that of the fastest decay curve of the CpcL-PBS complex, excited at 628 nm (Figure 1D). This suggests a 0.07-ns-scale energy transfer from PCB (F645) to red-PCB (F669). The second most rapid decay curve (*t*_c_ = 0.09 ns) featured large positive F669 and small F696 bands together with a large negative F729 band. This suggests a 0.09-ns-scale energy transfer from F669/F696 to F729. The pattern of the 0.09-ns curve is different from that of the 0.01-ns curve in Figure 1E, thus indicating a fast energy transfer from F696 to F729 inside PSI, which was detected upon the 405-nm excitation of the PBS-CpcL-PSI_4_ supercomplex (Figure 1E) or of PSI_4_ (Figure S6). Therefore, the slow 0.09-ns energy transfer from F669 to F696, but not the faster one from F696 to F729, seemed to limit the rate of overall energy transfer from CpcL-PBS (F669) to red-Chls (F729) in the supercomplex.

The 0.45-ns curve of the supercomplex with a positive peak at 729 nm (Figure 1F) resembled the 0.45–0.46-ns curve of PSI (Figure S6E), suggesting energy dissipation from F729. The 0.86-ns curve of the supercomplex, with positive peaks at 652 and 729 nm (Figure 1F), suggested energy dissipation from CpcL-PBS as F645 (the 0.6-ns curve in Figure 1D) and from PSI as F729 (the 0.77–0.78-ns curve in Figure S6E). The DAS of the supercomplex lacked long-lived components, like the 1.8-ns curve of CpcL-PBS, indicating the absence of free CpcL-PBS.

The short time constant of 0.09 ns for the energy transfer from CpcL-PBS-bound PCB to PSI_4_-bound Chls in the supercomplex was comparable to that of the energy transfer from APC-containing PBS to PSI/PSII.^8, 10, 17, 18^ Energy transfer from the rod-core-type PBS to PSII and PSI in the PBS-PSII-PSI megacomplex featured time constants of 0.08 and 0.15 ns, respectively.^8, 17, 18^ In this megacomplex, the slower energy transfer from PBS to PSI seems to be crucial for the optimally balanced PSII/PSI excitation to avoid photodamage.^19^ On the other hand, the fast energy transfer from CpcL-PBS to PSI via red-PCB in the PBS-CpcL-PSI_4_ supercomplex might be crucial for increasing the energy influx to PSI when higher PSI excitation is needed, e.g., under nitrogen starvation or green light conditions.^2, 12–14^

The efficiency of energy transfer (*Φ*) is defined as *Φ* = 1 - *τ*_fast_/*τ*, where *τ*_fast_ and *τ* are the fluorescence lifetimes of the donor, measured in the presence and absence of the acceptor, respectively. For the PBS-CpcL-PSI_4_ supercomplex, *Φ* was calculated as 95%, with *τ*_fast_ and *τ* at 0.09 (Figures 1F and 2) and 1.8 ns (Figure 1D and 2), respectively, according to energy transfer diagrams (Figure S8 and page S22-S24). The Förster radius for energy transfer from red-PCB to the Chl *a* acceptor was calculated as 6.5 nm using PhotochemCAD (pages S26–27).^20^ Accordingly, the 95% energy transfer efficiency of the supercomplex should be attained at distances of 3.2 and 3.9 nm with PCB and red-PCB, respectively, as the overlap integral (*J*-value in a ± 20-nm range from the emission peak) increases three-fold in the case of red-PCB (Figure S9). We assume that the binding of CpcL to PC contributes to the faster energy transfer to Chl *a* in PSI. This observation agrees with the prediction of a recent single-molecule experiment, claiming that the lower energy “red-PCB” state, which is induced by the specific interaction of PC with the linker protein CpcL (CpcG2), would lead to faster energy transfer to PSII/PSI in the absence of APC.^21^

Figure 3A shows a model proposed herein for energy transfer in the PBS-CpcL-PSI_4_ supercomplex. According to this model, light energy is transferred from PCB (F645) to red-PCB (F669) within the CpcL-PBS complex and to antenna Chl *a* in PSI_4_ with time constants of 0.07 and 0.09 ns, respectively, and is used by P700 for the photoreaction. The arrangements of PCB and Chl *a* molecules in the supercomplex were proposed by combining the predicted distance of 3.9 nm between red-PCB and Chls (Figures 3B and 3C) and the structure of the PBS-CpcL-PSI_4_ supercomplex, which was assumed by low-resolution single-particle EM.^2^ The CpcL-PBS moiety binds to two PSI monomers in PSI_4_, approximately 2 nm away from PsaC, PsaD, and PsaE subunits, and protrudes from the membrane surface toward the stroma. This structure was formulated to allow the docking of ferredoxin to the Fe-S cluster (F_B_) of PsaC in the PBS-CpcL-PSI_4_ supercomplex.^2^

**Figure 3.**
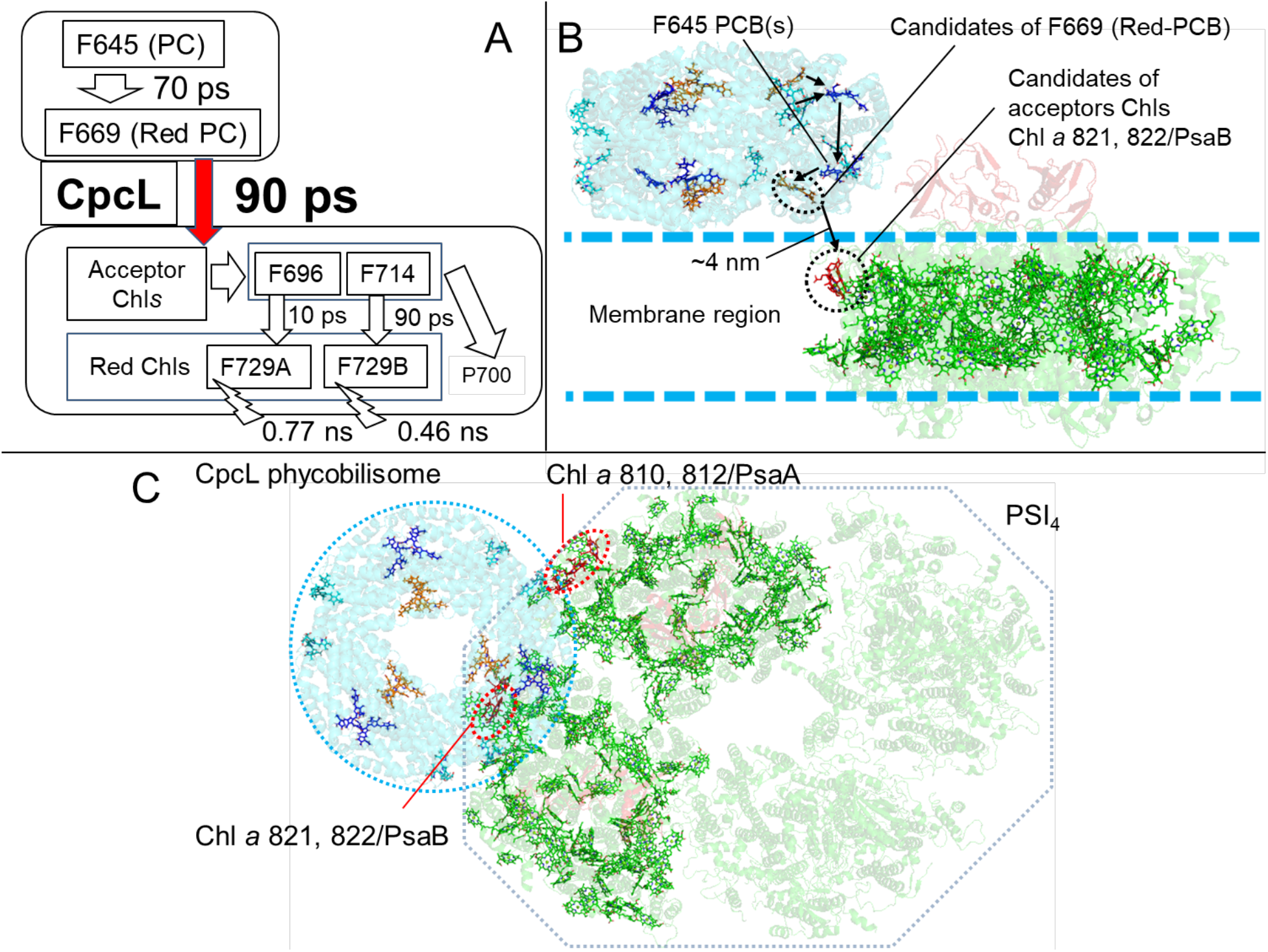
Energy transfer for the predicted arrangements of CpcL-PBS and PSI in the PBS-CpcL-PSI_4_ supercomplex. (A) Energy transfer pathways and time constants were estimated from the results of fluorescence analysis. The red arrow shows energy transfer from red-PCB on CpcL-PBS to the acceptor Chl *a*’s in PSI. Open arrows represent energy transfer directions. (B) Structure of the PBS-CpcL-PSI_4_ supercomplex estimated by single-particle EM analysis^2^ and the proposed pathways of energy transfer. Black arrows show excitation energy flows. For the structure modeling, the structures of PSI_4_ of *Anabaena* [PDB ID: 6JEO]^22^ and a PC hexamer from *Thermosynechococcus elongatus* [PDB ID: 5MJM]^22^ were used. (C) A view of the PBS-CpcL-PSI_4_ supercomplex from the stromal side. Aqua blue, blue, and orange denote PCB β-155, α-84, and β-84 in PC, respectively. Green denotes Chl *a* in PSI, while red denotes Chl *a* molecules that potentially accept energy from red-PCB.

The following pathway of energy transfer was proposed in the PBS-CpcL-PSI_4_ supercomplex: light energy absorbed by PCB β-155 is transferred to PCB β-84 and PCB α-84 in the same trimer, and then to PCB α-84 in the adjacent trimer but within the same hexamer (Figures 3B and 3C), as reported previously.^23^ It is likely that the 645-nm emission band of CpcL-PBS (Figure S3) originates from PCB α-84s/β-84s in the PC monomer moiety (Figures 3B and 3C) because the emission of PCB α-84s/β-84s occurs at 645 nm in a typical PCB.^23^ It is also assumed that the β-84 in the base of PC rod is the terminal emitter in the rod-core PBS that transfers energy to the APC core because the emission band of β-84 is red-shifted by the interaction with a linker domain of a rod-core linker CpcG.^24^ Similarly, β-84 can be a candidate for the red-PCB that emits F669 in CpcL-PBS. The structure model of the CpcL-PBS-PSI_4_ supercomplex implies that the nearest chlorophyll molecules to CpcL-PBS are Chls 821/822 in the PsaB of PSI, which were suggested to absorb at 670 nm.^25^ The estimated distance between PCB β-84 and Chls 821/822 (~4 nm) is short enough to allow the rapid and efficient energy transfer via the Förster mechanism. The red shifted emission band of PCB in CpcL-PCB gives the high spectral overlap with the absorption band of Chl *a* in PSI, and therefore, should be crucial for the efficient transfer of excitation energy from CpcL-PBS to PSI.

PCB α-84 at the base of PC rod can also be a candidate for the red-PCB in CpcL-PBS, judging by the estimated distance from Chls 821/822. Chls 810/812 in PsaA of a PSI monomer can be candidate acceptors by assuming the shift of binding angle of CpcL-PBS to ~30° along the membrane normal. The precise identification of donor and acceptor locations in PBS-CpcL-PSI_4_ supercomplexes will be accomplished by advanced structural analyses.

The present study measured the fast and direct energy transfer from CpcL-PBS to PSI_4_ in the PBS-CpcL-PSI_4_ supercomplex and proposed the corresponding energy transfer pathway (Figure 3). The binding of CpcL was estimated to induce an approximately 25-nm red shift of the absorption peak of PCB as the terminal emitter in the PC monomer. This red shift increases the overlap integral between the emission bands of PCB and the absorption bands of Chl *a* and extends the Förster radius. Consequently, fast and direct energy transfer with a 95% efficiency is achieved in the PBS-CpcL-PSI_4_ supercomplex over the 3.9-nm distance between Chl *a* on PSI and PCB in APC-free CpcL-PBS.

## EXPERIMENTAL METHODS

The PBS-CpcL-PSI_4_ supercomplex, PSI_4_, and CpcL-PBS were isolated from *Anabaena* sp. PCC 7120 as reported previously (pages S5–S6).^2^ The CpcL-PBS/PSI_4_ mixture was prepared as described on page S6. Time-resolved fluorescence profiles were obtained using a streak camera (#4334; Hamamatsu Photonics, Hamamatsu, Japan) and analyzed as reported previously (pages S7–S8).^15^ The diagrams of energy transfer kinetics and the rate equation solutions used to interpret the DASs of CpcL-PBS, PSI_4_, and the PBS-CpcL-PSI_4_ supercomplex are shown in Figures S8A–C and on pages S22–S24.

## ASSOCIATED CONTENT

The following files are available free of charge: sample preparation, absorbance, steady-state fluorescence, fitting results for fluorescence lifetime, discussion for control experiments, comparison with *T. elongatus* PSI trimer and *A. marina* PBS, estimation of rate constants, calculations according to the Förster theory.

## AUTHOR INFORMATION

### Notes

The authors declare no competing financial interests.

### Corresponding authors

1Present address: For Tomoyasu Noji; Research Center for Advanced Science and Technology, The University of Tokyo, 4-6-1 Komaba, Meguro-ku, Tokyo 153-8904, Japan. For Mai Watanabe; Institute of Biology III, Faculty of Biology, University of Freiburg, Freiburg, Germany

## ACKNOWLEDGMENT

The work was supported in part by the JST CREST program, and KAKENHI (Grant Numbers 17H05716. to M.W., 17K07440 and 20K06684 to S.I., and 17K18013 to T.N.). T.N. thanks the Takahashi Industrial and Economic Research Foundation and the Iwatani Naoji Foundation. The authors are grateful to Dr. Yoshimasa Fukushima for his helpful comments and discussions for the energy transfer.

## REFERENCES

(1) Watanabe, M.; Ikeuchi, M. Phycobilisome: Architecture of a Light-harvesting Supercomplex. Photosynth. Res. 2013, 116, 265–276.

(2) Watanabe, M.; Semchonok, D. A.; Webber-Birungi, M. T.; Ehira, S.; Kondo, K.; Narikawa, R.; Ohmori, M.; Boekema, E. J.; Ikeuchi, M. Attachment of Phycobilisomes in an Antenna-photosystem I Supercomplex of Cyanobacteria. Proc. Natl. Acad. Sci. U. S. A. 2014, 111, 2512–2517.

(3) Pizarro, S. A.; Sauer, K. Spectroscopic Study of the Light-harvesting Protein C-phycocyanin Associated with Colorless Linker Peptides. Photochem. Photobiol. 2001, 73, 556–563.

(4) Mullineaux, C. W. Excitation-energy Transfer from Phycobilisomes to Photosystem-I in a Cyanobacterium. Biochim. Biophys. Acta 1992, 1100, 285–292.

(5) Mullineaux, C. W. Excitation-energy Transfer from Phycobilisomes to Photosystem-I in a Cyanobacterial Mutant Lacking Photosystem-II. Biochim. Biophys. Acta 1994, 1184, 71–77.

(6) Kondo, K.; Ochiai, Y.; Katayama, M.; Ikeuchi, M. The Membrane-associated CpcG2-phycobilisome in *Synechocystis:* A New Photosystem I Antenna. Plant Physiol. 2007, 144, 1200–1210.

(7) Ueno, Y.; Aikawa, S.; Kondo, A.; Akimoto, S. Energy Transfer in Cyanobacteria and Red Algae: Confirmation of Spillover in Intact Megacomplexes of Phycobilisome and Both Photosystems. J. Phys. Chem. Lett. 2016, 7, 3567–3571.

(8) Liu, H. J.: Zhang, H.; Niedzwiedzki, D. M.; Prado, M.; He, G. N.; Gross, M. L.; Blankenship, R. E. Phycobilisomes Supply Excitations to Both Photosystems in a Megacomplex in Cyanobacteria. Science 2013, 342, 1104–1107.

(9) Nganou A. C.; David, L.; Adir, N.; Pouhe, D.; Deen, M. J.; Mkandawire, M. Evidence of Additional Excitation Energy Transfer Pathways in the Phycobiliprotein Antenna System of *Acaryochloris marina*. Photoch. Photobio. Sci. 2015, 14, 429–438.

(10) Theiss, C.; Schmitt, F. J.; Pieper, J.; Nganou, C.; Grehn, M.; Vitali, M.; Olliges, R.; Eichler, H. J.: Eckert, H. J. Excitation Energy Transfer in Intact Cells and in the Phycobiliprotein Antennae of the Chlorophyll *d* Containing Cyanobacterium *Acaryochloris marina*. J. Plant. Physiol. 2011, 168, 1473–1487.

(11) Watanabe, M.; Kubota, H.; Wada, H.; Narikawa, R.; Ikeuchi, M. Novel Supercomplex Organization of Photosystem I in *Anabaena* and *Cyanophora paradoxa*. Plant Cell Physiol. 2011, 52, 162–168.

(12) Hirose, Y.; Song, C. H.; Watanabe, M.; Yonekawa, C.; Murata, K.; Ikeuchi, M.; Eki, T. Diverse Chromatic Acclimation Processes Regulating Phycoerythrocyanin and Rod-Shaped Phycobilisome in Cyanobacteria. Mol. Plant 2019, 12, 715–725.

(13) Hirose, Y.; Shimada, T.; Narikawa, R.; Katayama, M.; Ikeuchi, M. Cyanobacteriochrome CcaS is the Green Light Receptor that Induces the Expression of Phycobilisome Linker Protein. Proc. Natl. Acad. Sci. U. S. A. 2008, 105, 9528–9533.

(14) Hirose, Y.; Narikawa, R.; Katayama, M.; Ikeuchi, M. Cyanobacteriochrome CcaS Regulates Phycoerythrin Accumulation in *Nostoc punctiforme*, a Group II Chromatic Adapter. Proc. Natl. Acad. Sci. U. S. A. 2010, 107, 8854–8859.

(15) Komura, M.; Shibata, Y.; Itoh, S. A New Fluorescence Band F689 in Photosystem II Revealed by Picosecond Analysis at 4-77 K: Function of Two Terminal Energy Sinks F689 and F695 in PSII. Biochim. Biophys. Acta 2006, 1757, 1657–1668.

(16) Shibata, Y.; Yamagishi, A.; Kawamoto, S.; Noji, T.; Itoh, S. Kinetically Distinct Three Red Chlorophylls in Photosystem I of *Thermosynechococcus elongatus* Revealed by Femtosecond Time-resolved Fluorescence Spectroscopy at 15 K. J. Phys. Chem. B 2010, 114, 2954–2963.

(17) Mullineaux, C. W.; Holzwarth, A. R. Kinetics of Excitation-energy Transfer in the Cyanobacterial Phycobilisome-Photosystem-II Complex. Biochim. Biophys. Acta 1991, 1098, 68–78.

(18) Mimuro, M.; Yamazaki, I.; Tamai, N.; Katoh, T. Excitation-energy Transfer in Phycobilisomes at −196°C Isolated from the Cyanobacterium *Anabaena variabilis* (M-3): Evidence for the Plural Transfer Pathways to the Terminal Emitters. Biochim. Biophys. Acta 1989, 973, 153–162.

(19) Yokono, M.; Takabayashi, A.; Akimoto, S.; Tanaka, A. A Megacomplex Composed of Both Photosystem Reaction Centres in Higher Plants. Nat. Commun. 2015, 6, 6675–6680.

(20) Dixon, J. M.; Taniguchi, M.; Lindsey, J. S. PhotochemCAD 2: A Refined Program with Accompanying Spectral Databases for Photochemical Calculations. Photochem. Photobiol. 2005, 81, 212–213.

(21) Gwizdala, M.; Kruger, T. P. J.; Wahadoszamen, M.; Gruber, J. M.; van Grondelle, R. Phycocyanin: One Complex, Two States, Two Functions. J. Phys. Chem. Lett. 2018, 9, 1365–1371.

(22) Meents, A.; Wiedorn, M. O.; Srajer, V.; Henning, R.; Sarrou, I.; Bergtholdt, J.; Barthelmess, M.; Reinke, P. Y. A.; Dierksmeyer, D.; Tolstikova, A.; Schaible, S.; Messerschmidt, M.; Ogata, C. M.; Kissick, D. J.; Taft, M. H.; Manstein, D. J.; Lieske, J.; Oberthuer, D.; Fischetti, R. F.; Chapman, H. N. Pink-beam Serial Crystallography. Nat. Commun. 2017, 8, 1281–1292

(23) Mimuro, M. Studies on Excitation-energy Flow in the Photosynthetic Pigment System - Structure and Energy-transfer Mechanism. Bot. Mag. Tokyo 1990, 103, 233–253.

(24) Ma, J. F.; You, X.; Sun, S.; Wang, X. X.; Qin, S.; Sui, S. F. Structural Basis of Energy Transfer in *Porphyridium purpureum* Phycobilisome. Nature 2020, 579, 146–151.

(25) Byrdin, M.; Jordan, P.; Krauss, N.; Fromme, P.; Stehlik, D.; Schlodder, E. Light Harvesting in Photosystem I: Modeling Based on the 2.5-Å Structure of Photosystem I from *Synechococcus elongatus*. Biophys. J. 2002, 83, 433–457.

