## supplemental file for "Direct Energy Transfer from Allophycocyanin-Free Rod-Type CpcL-Phycobilisome to Photosystem I"

[d]JSPS Overseas Research Fellow

[e]Department of Physics, Graduate School of Nagoya University, Furo-cho, Chikusa-ku Nagoya, Aichi 464-8602, Japan

AUTHOR INFORMATION

**Corresponding Author**

*Tomoyasu Noji, [1]Research Center for Advanced Science and Technology, The University of Tokyo, Tokyo 153-8904, Japan, Tel.: +81-6-6605-3486,

*Mai Watanabe, Institute of Biology III, Faculty of Biology, University of Freiburg, Freiburg, Germany, Tel.: +49-761-2036948,

### Contents

**Materials and Methods (pages S5–S8)**

***Sample preparation*** **(page S5–S6)**

***Measurement and analysis of time-resolved fluorescence*** **(pages S7–S8)**

***Measurement of steady-state fluorescence*** **(page S8)**

***Measurement of absorption spectra*** **(page S8–S9)**

**Results and discussion (pages S10–S29)**

***Isolation of $PSI_4$, PBS-CpcL-$PSI_4$ supercomplex, and CpcL-PBS*** **(page S10)**

**Figure S1.** (A) Isolation of $PSI_4$, PBS-CpcL-$PSI_4$ supercomplex, and CpcL-PBS by sucrose gradient centrifugation. (B) Absorption spectra of $PSI_4$, PBS-CpcL-$PSI_4$ supercomplex, CpcL-PBS, and the CpcL-PBS/$PSI_4$ mixture. (page S11)

**Figure S2.** Absorption spectrum of CpcL-PBS and the second derivative spectrum. (page S12)

***Steady-state absorbance/fluorescence/excitation spectra of $PSI_4$, PBS-CpcL-$PSI_4$ supercomplex, and CpcL-PBS at 77 K*** (pages S13–S14)

**Figure S3.** (A) Fluorescence spectra of CpcL-PBS and PBS-CpcL-$PSI_4$ supercomplex measured at 77 K. Absorption spectrum of PBS-CpcL-$PSI_4$ supercomplex at 77 K. Excitation spectrum of CpcL-PBS alone monitored at 728 nm. (B) Comparison of excitation spectra of PBS-CpcL-PSI$_4$ supercomplex (red) and PSI$_4$ (black) at 77 K. Emission wavelength is 729 nm. (page S15)

**Figure S4.** Comparison of absorption or fluorescence spectra of *Anabaena* PBS-CpcL-PSI$_4$ supercomplex and *Thermosynechococcus elongatus* PSI trimer at 77K. (page S16)

***Fluorescence lifetime at 77K*** **(pages S17–S18)**

**Figure S5.** Fluorescence lifetime values and global fitting results of the samples. (page S19)

**Figure S6.** Two-dimensional fluorescence images measured at 77 K and DASs (with time constants) calculated from these images of PSI4 and the CpcL-PBS/PSI$_4$ mixture. (page S20)

**Figure S7.** Time-resolved fluorescence spectra at 77 K of the CpcL-PBS/PSI$_4$ mixture and PBS-CpcL-PSI$_4$ supercomplex excited at 628 nm. (page S21)

***Estimation of the rate constants*** **(pages S22–S24)**

**Figure S8.** Energy transfer diagrams for CpcL-PBS, PSI$_4$, and PBS-CpcL-PSI$_4$ supercomplex. (page S25)

***Calculations according to the Förster theory*** **(page S26–S27)**

**Figure S9.** Overlap integral of fluorescence and absorbance. (page S27)

**References (pages S28–S29)**

#### Materials and Methods

##### *Sample preparation*

*Anabaena* sp. PCC 7120 cells were grown at 31 °C in liquid BG11 supplemented with 20 mM Hepes·potassium hydroxide (pH 8.2; Rippka R (1988) Isolation and purification of cyanobacteria. Methods Enzymol 167:3–27.) and with bubbling of 1% $CO_2$ under continuous illumination with white fluorescent lamps (ca. 20 $\mu$mol photons $m^{-2}\cdot s^{-1}$).

PSI complexes were isolated as previously described.[1] The thylakoid membranes isolated from vegetative cells of the cyanobacterium *Anabaena* sp. PCC 7120 at a concentration equivalent to 1 mg Chl/mL were solubilized with 1% n-dodecyl-β-D-maltoside (DDM) for 30 min on ice, followed by centrifugation at 300,000 × *g* for 30 min at 4°C. The obtained supernatant was diluted with three volumes of a buffer containing 50 mM 4-(morpholino) ethanesulfonic acid·sodium hydroxide (MES-NaOH; pH 6.5), 10 mM $MgCl_2$, and 5 mM $CaCl_2$, loaded onto a 10%–30% linear sucrose density gradient in the same solution but with DDM, and centrifuged at 130,000 × *g* for 6 h at 4°C to obtain the purified $PSI_4$ and PBS-CpcL-$PSI_4$ supercomplex.

PBS was isolated as previously described,[1] with some modifications. Briefly, cells grown in BG11 were harvested, washed twice with 0.9 M potassium phosphate buffer (pH 7.0), and lysed with zirconia beads using a bead beater (Micro Smash MS-100R; Tomy Digital Biology, Tokyo, Japan). The cell extract was treated with 2% (v/v) Triton X-100 in 0.9 M phosphate buffer for 30 min and then centrifuged at 20,000 × *g* and 18°C for 20 min to separate the mixture into an upper green Triton X-100 layer and a lower blue aqueous layer. The latter was loaded onto a 10%–50% (w/v) linear sucrose density gradient with 0.9 M phosphate buffer and centrifuged at 130,000 × *g* for 16 h at 18°C to obtain purified CpcL-PBS.

Concentrations of samples were adjusted on ice just after purification for spectroscopic analysis. The concentrations of $PSI_4$, PBS-CpcL-$PSI_4$ supercomplex, and CpcL-PBS/$PSI_4$ mixture were adjusted to an optical density at 680 nm ($OD_{680}$) of 0.05–0.06 for a 1.0-mm light path. Purified $PSI_4$ and PBS-CpcL-$PSI_4$ supercomplex were suspended in a low-salt buffer [50 mM MES (pH 6.5), 10 mM $MgCl_2$, 5 mM $CaCl_2$, 10% (w/v) sucrose]. To make the CpcL-PBS/PSI mixture, CpcL-PBS, $PSI_4$, high-salt buffer [0.9M potassium phosphate (pH 7.0), 10% (w/v) sucrose], and low-salt buffer were mixed to a 1:1 stoichiometric ratio of CpcL-PBS to $PSI_4$ mixture (the ratio of OD630/OD680 = 0.3 in the final mixture, so it was equivalent to the ratio in PBS-CpcL-$PSI_4$ supercomplex), and the ratio of high-salt buffer and low-salt buffer to 4:1. The concentration of the CpcL-PBS solution was adjusted to $OD_{630}$ = 0.03 with a light path of 1.0 mm by using high-salt buffer. For the measurement of time-resolved fluorescence, all samples were put in a sample holder and quickly frozen by liquid $N_2$. The samples were stored at -80°C until measurement.

***Measurement and analysis of time-resolved fluorescence***

Time-resolved fluorescence profiles were obtained on a spectrograph consisting of a 30-cm monochromator and streak camera (#4334; Hamamatsu Photonics, Hamamatsu, Japan). All samples were immediately put in a liquid nitrogen cryostat to measure fluorescence lifetime at 77K before thawing of samples. The sample in a cuvette with a light path of 1 mm was cooled to 77 K in liquid nitrogen and then placed in a liquid nitrogen cryostat (Oxford Instruments, Abingdon, Oxfordshire, UK). Excitation was provided by a laser diode emitting at a wavelength of 405 or 628 nm, operating at 1 MHz with a 50-ps pulse duration (Hamamatsu Photonics). For measurements with 628-nm excitation, a 630-nm interference filter was placed in the light path to eliminate stray light from the laser. Fluorescence from the sample was focused onto the entrance slit of a monochromator with a grating of 100 grooves/mm (Chromex 2501-S; Hamamatsu Photonics) and a slit width of 50 μm through a 650-nm long-pass filter (TS OD2; Edomund, Tokyo, Japan). The monochromator was set at a center wavelength of 705 nm with a spectral resolution of 4.1 nm. The beam exiting the monochromator was focused onto the streak camera to divide photoelectrons according to their arrival times, and traces of photoelectrons on the lucifer plate of the streak output were collected at a resolution of 640 × 480 pixels with a charge-coupled device (CCD) camera. Each trace was analyzed in the photon-counting mode to yield a typical 150-nm wavelength width (x-axis) and 2- or 5-ns duration (y-axis) for 100,000 shots (approximately 1 h).[19] The sensitivity of the CCD image was corrected using a standard light source (SL1-CAL; StellarNet, Tampa, FL, USA) and a spectrometer (EPP2000-UVN-SR; StellarNet). The measured fluorescence profiles were divided into 16 wavelength regions (40 pixels of 8.8 nm each) and summed along the wavelength axis to obtain the fluorescence time course in each region. Global multi-exponential fitting was performed for these time courses by convolution to obtain the decay-associated spectra (DAS).[20] A Gaussian shape was assumed as a response function.

***Measurement of steady-state fluorescence***

Steady-state fluorescence spectra were obtained at 77 K with a fluorescence spectrophotometer (FluoroMax-4; Horiba, Kyoto, Japan) equipped with a home-built Dewar vessel containing liquid nitrogen. Samples were transferred to a sample cuvette with a light path of 2 mm. The excitation wavelength for fluorescence measurements was 440 or 600 nm. Fluorescence from the sample was detected through a 620-nm red cut-off filter (R-62; Hoya Candeo Optronics, Saitama, Japan) at 1-nm resolution. Buffer conditions of samples were the same as fluorescence lifetime measurements and absorption spectra measurements at room temperature.

***Measurement of absorption spectra***

Absorption spectra were measured at room temperature with a spectrophotometer (UV-1800PC; Shimadzu, Kyoto, Japan). Samples were prepared as shown in sample preparation section above.

The absorption spectrum of the PBS-CpcL-$PSI_4$ supercomplex was measured at 77 K with a spectrophotometer (UV-3100PC; Shimadzu). The concentration was adjusted to 4 μg Chl/mL in a buffer composed of 7 mM MES-NaOH (pH 6.5), 3 mM $MgCl_2$, 2 mM $CaCl_2$,

3% (w/v) sucrose, 0.003% (w/v) DDM, and 66% (v/v) glycerol. The sample was transferred to a cuvette with a light path of 10 mm, cooled to 77 K in liquid nitrogen, and placed in a liquid nitrogen cryostat (Oxford Instruments, Abingdon, Oxfordshire, UK).

#### Results and discussion

##### *Isolation of $PSI_4$, PBS-CpcL-$PSI_4$ supercomplex, and CpcL-PBS*

$PSI_4$ and PBS-CpcL-$PSI_4$ supercomplex were isolated from the cyanobacterium *Anabaena* sp. PCC 7120 using a linear sucrose density gradient containing a low-salt buffer and 0.01% DDM (Figure S1A, lane 1). CpcL-PBS was fractionated using the linear sucrose density gradient in the presence of high concentrations of phosphate (Figure S1A, lane 2). Absorption spectra of the isolated $PSI_4$, PBS-CpcL-$PSI_4$ supercomplex, CpcL-PBS, and a mixture of CpcL-PBS and $PSI_4$ (CpcL-PBS/$PSI_4$ mixture) at room temperature are shown after normalization at the peak with the longest wavelength (Figure S1B). The ratio of $OD_{630}/OD_{680}$ of the CpcL-PBS/$PSI_4$ mixture was adjusted to 0.30 to meet the ratio in the purified PBS-CpcL-$PSI_4$ supercomplex, in which CpcL-PBS and $PSI_4$ were approximately 1:1.[1]

The second derivative of the absorbance spectrum of CpcL-PBS at room temperature revealed a band at 635 nm (Figure S2), which differed from the 641-nm peak of the PC/AP heterohexamer in *Acaryochloris marina* estimated using a similar method.[2] Therefore, the structure of PCB in CpcL-PBS responsible for binding to $PSI_4$ is different from that in the ordinary PC/AP heterohexamer complex that binds to PSII.

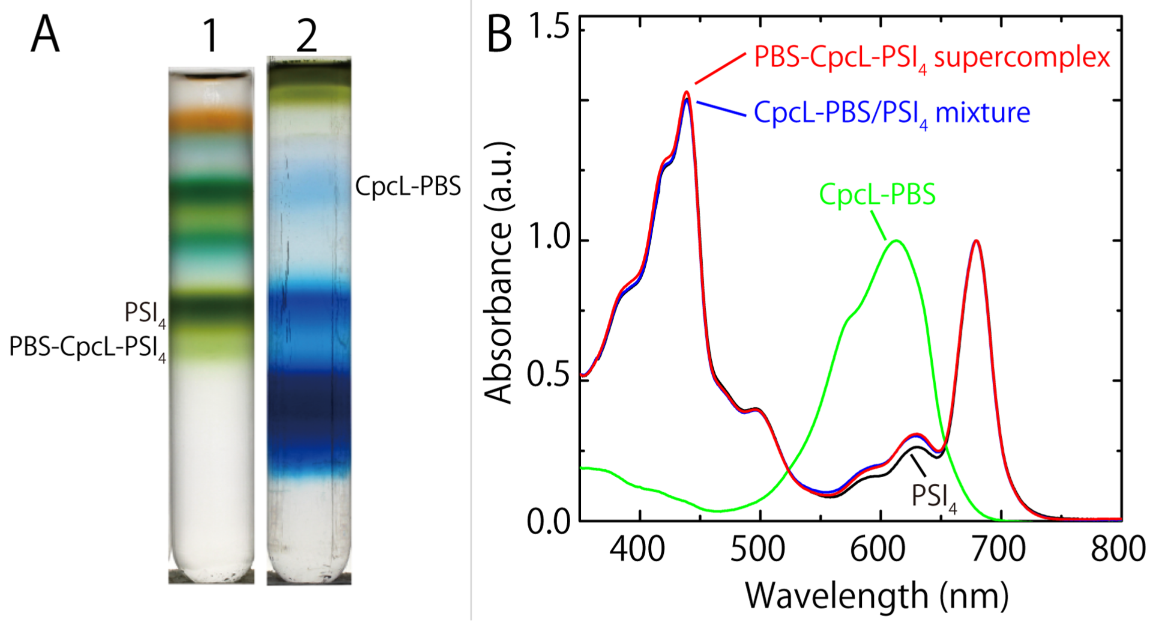


**Figure S1.** (A) Isolation of $PSI_4$, PBS-CpcL-$PSI_4$ supercomplex, and CpcL-PBS by sucrose gradient centrifugation. Lane 1 shows the separation of complexes from the thylakoid after centrifugation through a linear sucrose gradient containing low-salt buffer and 0.01% (w/v) DDM. Lane 2 shows the separation of CpcL-PBS in the presence of high concentrations of phosphate. (B) Absorption spectra of $PSI_4$ (black), PBS-CpcL-$PSI_4$ supercomplex (red), CpcL-PBS (green), and the CpcL-PBS/$PSI_4$ mixture (blue) at room temperature, normalized to the peak of the longest wavelength. Detailed description of the results is provided page S10.

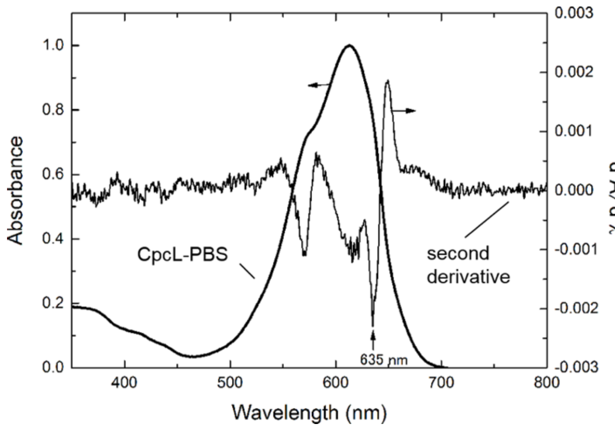


**Figure S2.** Absorption spectrum of CpcL-PBS (thick line) and its second derivative spectrum (thin line). Detailed description of the results is provided on page S10.

***Steady-state absorbance/fluorescence/excitation spectra of $PSI_4$, PBS-CpcL-$PSI_4$ supercomplex, and CpcL-PBS at 77 K***

The excitation spectrum for the 728-nm emission peak of CpcL-PBS at 77 K revealed peaks at 568, 614, and 633 nm (Figure S3A). A small shoulder at 659 nm suggested the presence of a terminal emitter(s). Hence, the bands at 614 and 633 nm were assigned to originate from PCBs in PC.[1] The band at 659 nm has not been observed in typical PCs at 77 K[3] and was suggested to originate from the red-shifted PCB(s) in CpcL-PBS.

The fluorescence spectrum of CpcL-PBS, excited at 600 nm at 77 K showed two major peaks at 645 and 669 nm (Figure S3A), suggesting that they were emitted from the absorption bands at 633 and 659 nm, respectively. In contrast, typical PC hexamers with a CpcC rod linker and a CpcG rod–core linker are reported to give absorbance peaks at 643 and 652 nm at 77 K, respectively,[4] and the AP cores are known to emit at 685 nm.[4,5] Hence, it is likely that the emission band at 645 nm (F645) of CpcL-PBS originated from PCB bound to a typical PC, whereas that at 669 nm (F669) derived from PCB bound to PC that interacted with CpcL. Therefore, we named the 669-nm terminal emitter “red PCB”.

The absorption spectrum of the PBS-CpcL-$PSI_4$ supercomplex overlapped with the fluorescence spectrum of CpcL-PBS at 660–680 nm (Figure S3A). This range overlaps with the Qy absorption band of Chl *a* of the core antenna in $PSI_4$. The absorption peaks of PBS-CpcL-$PSI_4$ supercomplex were at 679 and 684 nm with shoulders near 670 nm (Figure S3A). The fluorescence intensity of the PBS-CpcL-$PSI_4$ supercomplex excited at 550-650 nm was higher than that of $PSI_4$ (Figure S3B). Chl absorption bands at 670 nm in the core antenna in $PSI_4$ will, therefore, efficiently accept excitation energy from the F669 emission band of terminal emitters in CpcL-PBS. The 670-nm absorbing Chl *a* molecules may be located on the exterior of PSI, where the latter accepts excitation energy from red PCBs, and then transfers it to P700 as well as to the 679-nm and 684-nm absorbing Chls.[6]

TEM analysis of PBS-CpcL-$PSI_4$ supercomplex suggests that CpcL-PBS is located on the external surface of $PSI_4$ and is rather far from P700 in the central moiety.[1] Thus, energy transfer from CpcL-PBS to P700 may be achieved via Chls located on the exterior of $PSI_4$. The peaks of the Qy absorption band of Chl *a* molecules in the PBS-CpcL-$PSI_4$ supercomplex differed from those in the PSI trimer ($PSI_3$) from *Thermosynechococcus elongatus* (Figure S4A). The results suggest that the structure of the core antenna moiety of $PSI_4$ differs slightly from that of *T. elongatus* $PSI_3$, even though the overall structure of the PSI monomer appears to be very similar between the two species.[1]

The peak and shoulder bands at approximately 710 and 720 nm in the absorption spectrum of PBS-CpcL-$PSI_4$ supercomplex indicate the presence of red-Chls, similar to the one in $PSI_3$ from *T. elongatus* (Figure S4A). The emission peak of PBS-CpcL-$PSI_4$ supercomplex excited at 440 nm at 77 K was observed at 729 nm (F729), as was that of *T. elongatus* $PSI_3$ (Figure S4B). These results suggest that the structure around red Chls in $PSI_4$ is similar to that in *T. elongatus* $PSI_3$. The amino acid sequences of PsaA/B in *Anabaena* PSI are almost identical to those in *T. elongatus* PSI.[1]

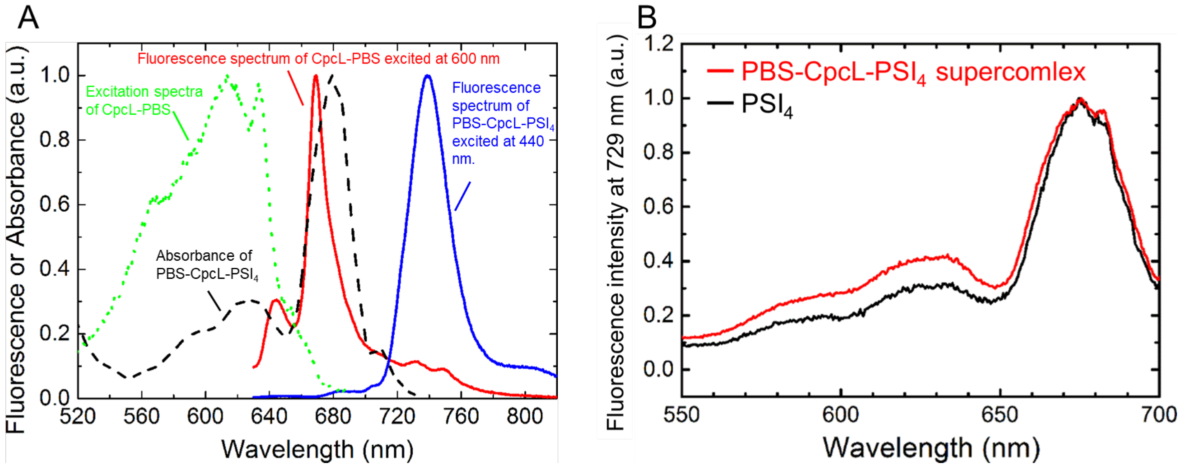


**Figure S3.** (A) Fluorescence spectra of CpcL-PBS (red line) and the PBS-CpcL-$PSI_4$ supercomplex (blue line) measured at 77 K. CpcL-PBS and the PBS-CpcL-$PSI_4$ supercomplex were excited at 600 and 440 nm, respectively. The broken line shows the absorption spectrum of the PBS-CpcL-$PSI_4$ supercomplex at 77 K, whereas the green dotted line shows the excitation spectrum of CpcL-PBS alone at 728 nm. Detailed description of the results is provided on pages S13–S14. (B) Comparison of excitation spectra of the PBS-CpcL-$PSI_4$ supercomplex (red) and $PSI_4$ (black) at 77 K. Emission wavelength is 729 nm. Detailed description of the results is provided on main text.

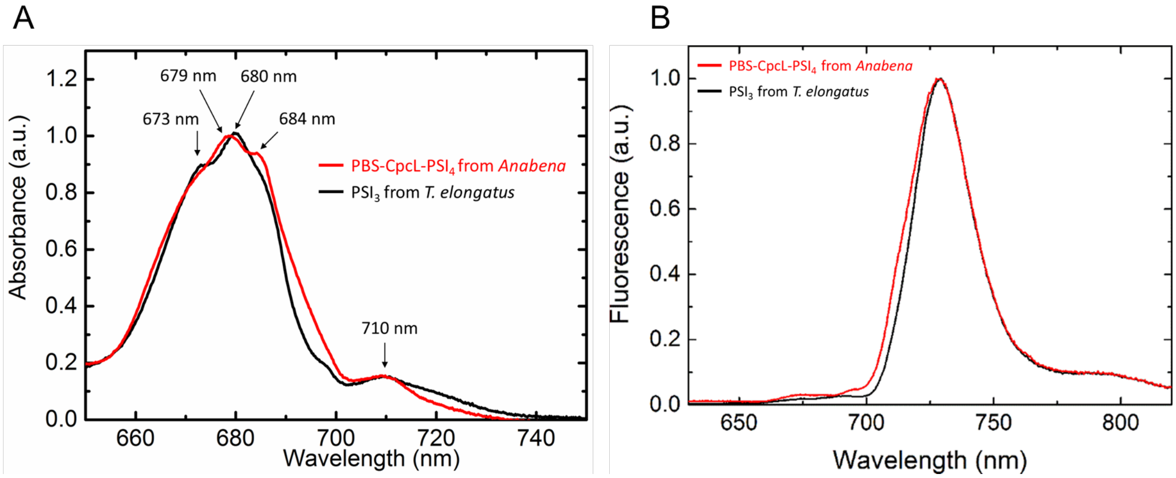


**Figure S4.** Comparison of absorption (A) and fluorescence (B) spectra of *Anabaena* PBS-CpcL-$PSI_4$ supercomplex (red) and *T*. *elongatus* $PSI_3$ (black) at 77 K. The excitation wavelength is 440 nm. Detailed description of the results is provided on page S13–S14.

***Fluorescence lifetime at 77 K***

Figure S5 shows global fitting results of the samples.

Figure S6 shows the wavelength-time two-dimensional (2D) images of fluorescence emission of the samples at 77 K. The main emission band at 729 nm of $PSI_4$ excited at 405 nm (Figure S6C) was assigned to F729 (Figure S3). Band intensity was lower in $PSI_4$ excited at 628 nm (Figure S6A) than in $PSI_4$ excited at 405 nm due to lower absorption at 628 nm. The similarity between Figures S6A and S6C suggests similar energy transfer processes inside $PSI_4$ even after excitation at different wavelengths.

The wavelength-time 2D image of the CpcL-PBS/$PSI_4$ mixture (Figure S6B) was similar to that obtained by superposing Figures S6A and 1A, suggesting no energy transfer between CpcL-PBS and $PSI_4$.

Figure S6E shows the decay-associated spectra (DAS) of $PSI_4$ from the fluorescence image excited at 405 nm at 77 K (Figure S6C). A DAS component of ~0.01 ns suggested energy transfer from the 696-nm Chl *a* band (positive band, F696) to red-Chls at 729 nm (negative band, named F729A; Figure S6E). The component at 0.09 ns showed energy transfer from a 714-nm band (F714) to a 729-nm band (named F729B), whereas components at 0.45 and 0.78 ns suggested thermal dissipation of excitation energy from red Chls at 729 nm (Figure S6E). A DAS component of 2.2 ns, whose time constant was comparable to the natural lifetime of Chl *a*, was small.

The fluorescence kinetics of the CpcL-PBS/$PSI_4$ mixture and PBS-CpcL-$PSI_4$ supercomplex were analyzed globally by assuming 3–4 multi-exponential components. The DAS of the CpcL-PBS/$PSI_4$ mixture excited at 628 nm revealed a 0.04-ns DAS (Figure S6D) that could be interpreted as a mixture of the 0.01-ns DAS of $PSI_4$ shown in Figure S6E and the 0.07-ns DAS of CpcL-PBS shown in Figure 1D. This provides evidence for two independent pathways of energy transfer; one from F645 to F669 in CpcL-PBS, and the other from bulk antenna Chls in PSI to red Chls in $PSI_4$. The 0.38-ns DAS in Figure S6D resembled the 0.45-ns DAS in Figure S6E, indicating energy dissipation in red Chls. The 0.69-ns DAS in Figure S6D appears to represent the mixture of the 0.78-ns DAS of $PSI_4$ shown in Figure S6E and the 0.6-ns DAS of CpcL-PBS shown in Figure 1D. The 1.72-ns DAS in Figure S6D resembled the 1.8-ns DAS of CpcL-PBS shown in Figure 1D, suggesting that there was no energy transfer between CpcL-PBS and $PSI_4$ in the mixture.

Figure S7 shows the time-resolved fluorescence spectra of the CpcL-PBS/$PSI_4$ mixture and the PBS-CpcL-$PSI_4$ supercomplex excited at 628 nm. The decrease in F669 intensity and the increase in F729 intensity suggest efficient energy transfer from CpcL-PBS to $PSI_4$ in PBS-CpcL-$PSI_4$ supercomplex (Figure S7B).

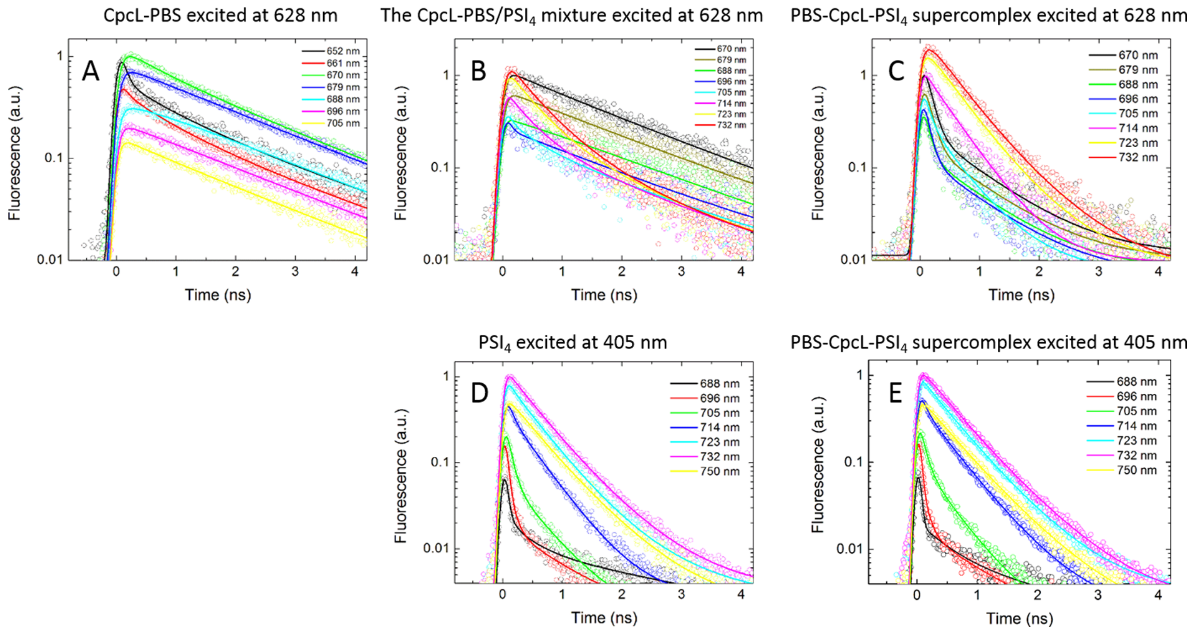


**Figure S5.** Fluorescence lifetimes and global fitting results of the four samples. Open circles and traces show raw data and fitted results, respectively. CpcL-PBS (A), the CpcL-PBS/$PSI_4$ mixture (B), and PBS-CpcL-$PSI_4$ supercomplex (C) excited at 628 nm; $PSI_4$ (D), and PBS-CpcL-$PSI_4$ supercomplex (E) excited at 405 nm.

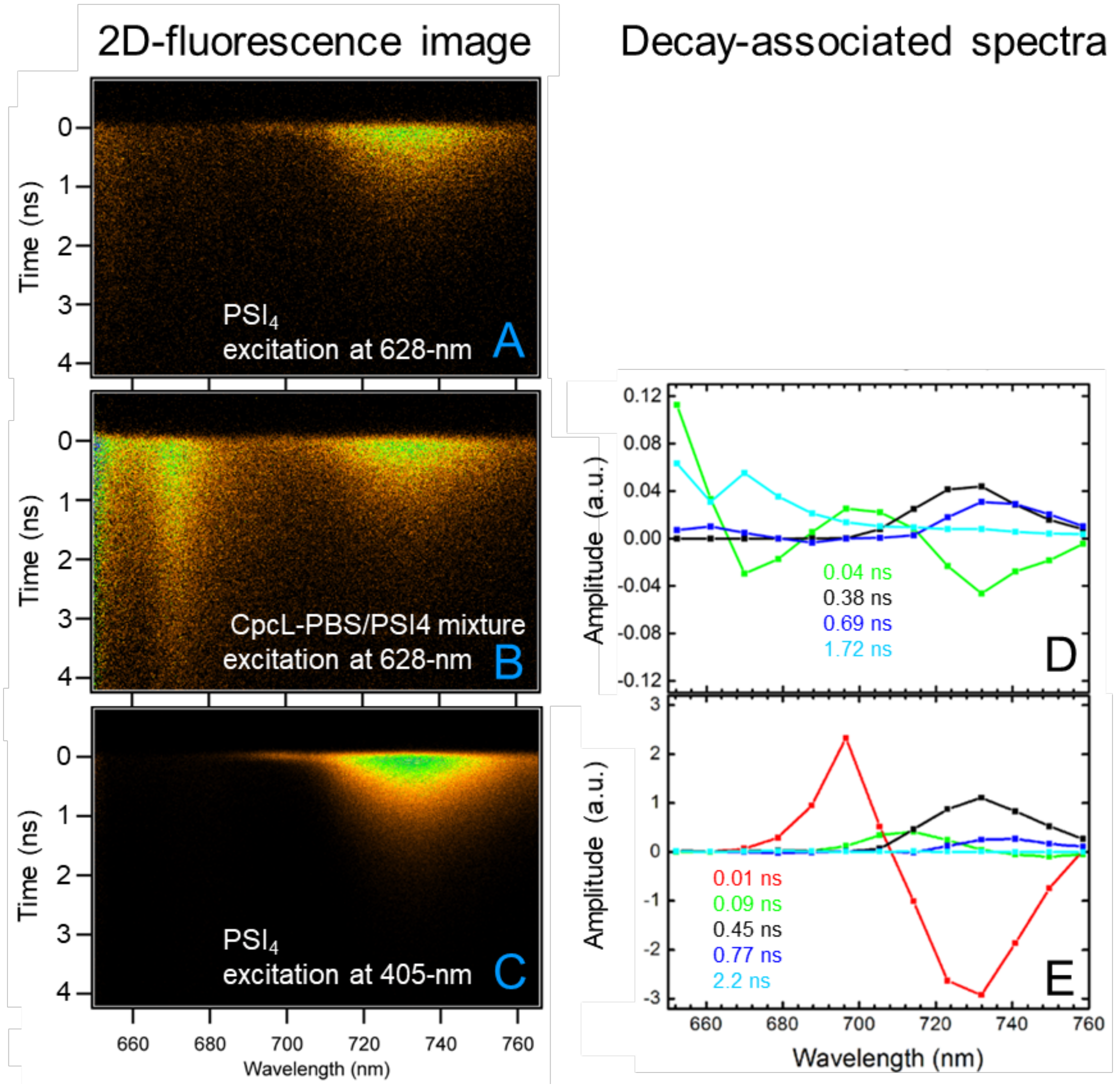


**Figure S6.** Two-dimensional fluorescence images measured at 77 K (A–C), and DASs (with time constants) calculated from these images (D–E). (A) $PSI_4$ excited at 628 nm. (B, D) CpcL-PBS/$PSI_4$ mixture excited at 628 nm. (C, E) $PSI_4$ excited at 405 nm. Detailed description of the results is provided on pages S17–S18.

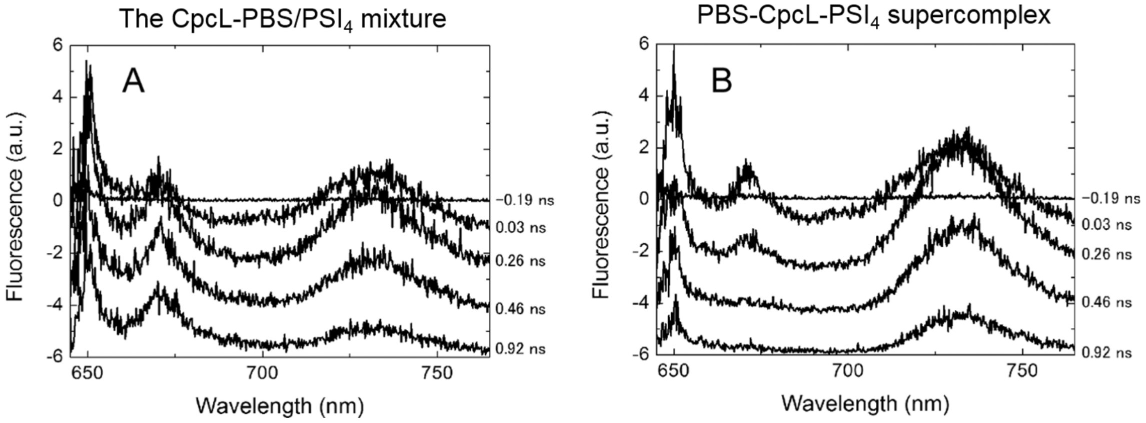


**Figure S7.** Time-resolved fluorescence spectra at 77 K of the CpcL-PBS/$PSI_4$ mixture (A) and PBS-CpcL-$PSI_4$ supercomplex (B) excited at 628 nm. Detailed description of the results is provided on pages S17–S18.

***Estimation of the rate constants in the energy transfer diagram for the CpcL-PBS complex***

A simple energy transfer diagram of CpcL-PBS was assumed, as shown in Figure S8A, by ignoring the minor 0.6-ns DAS. The equations (eq.) (1) and (2) below give the solution for the diagram in Figure S8A. Where, $k_1$-$k_8$ are rate constants that are defined in Figure S8, $t$ is time, and $A_1$-$A_4$ are amplitude.

$$[F645] = A_1 \exp(-(k_1 + k_2)t) \qquad \text{eq. 1}$$

$$[F669] = A_2 \exp(-(k_1 + k_2)t) + A_3 \exp(-k_3 t) \qquad \text{eq. 2}$$

By assuming that the long lifetime DAS in Figure 1D represents intrinsic lifetimes of F645 and F669, $k_1$ and $k_3$ can be expressed as below.

$$k_1 = k_3 = (1.8\, ns)^{-1} \qquad \text{eq. 3}$$

We can assume that the fastest DAS in Figure 1D represents the energy transfer from F645 to F669, so that $k_1 + k_2$ can be expressed as below.

$$k_1 + k_2 = (0.07\, ns)^{-1} \qquad \text{eq. 4}$$

From eqs. (3) and (4), the rate constant of energy transfer $(k_2)$ was calculated as below.

$$k_2 = (0.07\,ns)^{-1} - (1.8\,ns)^{-1} = (0.073\,ns)^{-1} \cong (0.07\,ns)^{-1} \qquad \text{eq. 5}$$

***Estimation of the rate constants in the energy transfer diagram for PSI$_4$***

Energy transfer from bulk Chls to F696 or F714 should be faster than the time resolution of the present streak camera system. Therefore, the energy transfer diagram in PSI$_4$ was assumed as Figure S8B and was used to interpret the DAS in Figure S6E. The solution of the rate equations developed from Figure S8B is presented in eqs. (6–9).

$$[F696] = A_1 \exp(-k_5 t) \qquad \text{eq. 6}$$

$$[F729A] = A_2 \exp(-k_5 t) + A_3 \exp(-k_6 t) \qquad \text{eq. 7}$$

$$[F714] = A_3 \exp(-k_7 t) \qquad \text{eq. 8}$$

$$[F729B] = A_4 \exp(-k_7 t) + A_3 \exp(-k_8 t) \qquad \text{eq. 9}$$

DAS values of 10 ps, 0.45 ns, 90 ps, and 0.78 ns in Figure S6E are assumed to represent the energy transfer from F696 to F729A, the quenching in F729A, the energy transfer from F714 to F729B, and the quenching in F729B, respectively. The rate constants $k_5$, $k_6$, $k_7$, and $k_8$ are determined as below, respectively.

$$k_5 = (0.01\,ns)^{-1}, k_6 = (0.45\,ns)^{-1}, k_7 = (0.09\,ns)^{-1}, k_8 = (0.78\,ns)^{-1} \qquad \text{eq. 10}$$

***Estimation of the rate constants in the energy transfer diagram for the rod-PBS-CpcL-$PSI_4$ supercomplex***

A simple energy transfer diagram for the CpcL-PBS-$PSI_4$ supercomplex was assumed as shown in Figure S8C. The solution of the rate equation developed from Figure S8C gives eqs. (11) and (12).

$$[F645] = A_1 \exp(-(k_1 + k_2)t) \qquad \text{eq. 11}$$

$$[F669] = A_2 \exp(-(k_1 + k_2)t) + A_3 \exp(-(k_3 + k_4)t) \qquad \text{eq. 12}$$

The rate constants $k_1$, $k_2$, and $k_3$ are determined from the equations (3) and (4) as below, respectively.

$$k_1 = k_3 = (1.8\,ns)^{-1}, \qquad k_1 + k_2 = (0.07\,ns)^{-1}$$

Because the DAS corresponding to the first lifetime in Figure 1F indicates the energy transfer from F669 to PSI, $k_3 + k_4$ are determined as below.

$$k_3 + k_4 = (0.09\,ns)^{-1} \qquad \text{eq. 13}$$

Based on equations (3), (4), and (13), the rate constant of energy transfer $(k_4)$ is calculated as below.

$$k_4 = (0.09\,ns)^{-1} - (1.8\,ns)^{-1} = (0.094\,ns)^{-1} \cong (0.09\,ns)^{-1} \qquad \text{eq. 14}$$

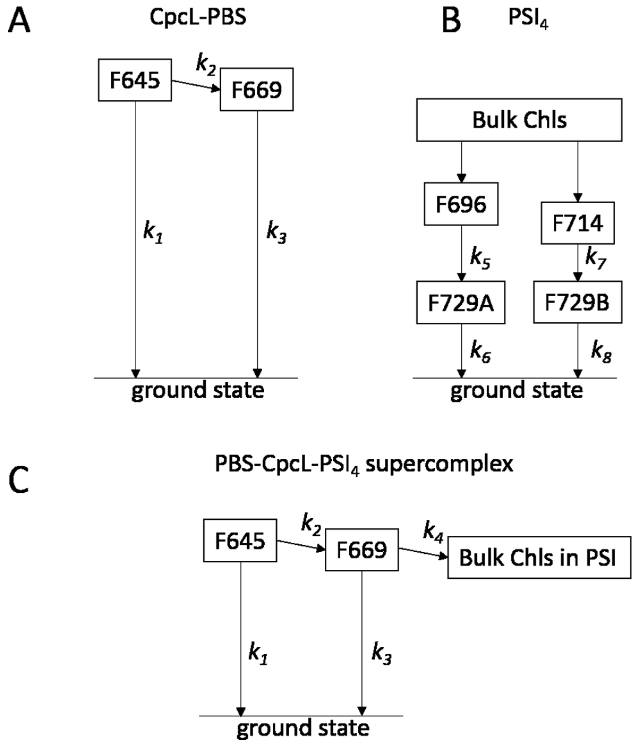


**Figure S8.** Energy transfer diagrams for CpcL-PBS (A), $PSI_4$ (B), and the PBS-CpcL-$PSI_4$ supercomplex (C).

***Calculations according to the Förster theory***

The efficiency ($\Phi$) of energy transfer between CpcL-PBS and $PSI_4$ was calculated using PhotochemCAD.[7] The relationship between $\Phi$ and the distance ($R$) between a donor and an acceptor is described by eq. (15).

$$\Phi = R_0^6 / (R^6 + R_0^6) \qquad \text{eq.15}$$

The Förster radius ($R_0$) is defined by eq. (16).

$$R_0^6 = (8.8 \times 10^{23})\ \kappa^2\ \Phi_f\ J\ n^{-4} \qquad \text{eq.16}$$

$$J = \int f_D(\lambda)\varepsilon_A(\lambda)\lambda^4 d\lambda \qquad \text{eq. 17}$$

In eq. 17, $f_D$ and $\varepsilon_A$ are the fluorescence spectrum of a donor and absorption spectrum of an acceptor, respectively. The orientation factor ($\kappa^2$) was determined to be 2/3, as in the case of random orientation. The fluorescence yield of the donor ($\Phi_f$) was determined to be 0.48 according to Ref. 8. $J$ is the spectral overlap integral (range of 649–689 nm) and was calculated from the emission spectrum of the donor (CpcL-PBS) and absorption spectrum of an acceptor Chl in the core antenna within PSI. The red trace in Figure S3 was used as the fluorescence spectrum of CpcL-PBS. The absorption spectrum of the acceptor Chl was assumed to be similar to that in the spectrum of PBS-CpcL-$PSI_4$ supercomplex (black broken line in Figure S3), and its absorption coefficient was assumed to be 79290 $M^{-1}$ $cm^{-1}$ (i.e., for the value in N,N'-dimethylformamide).[9] The refraction index ($n$) was 1.333 (i.e., the value for water).

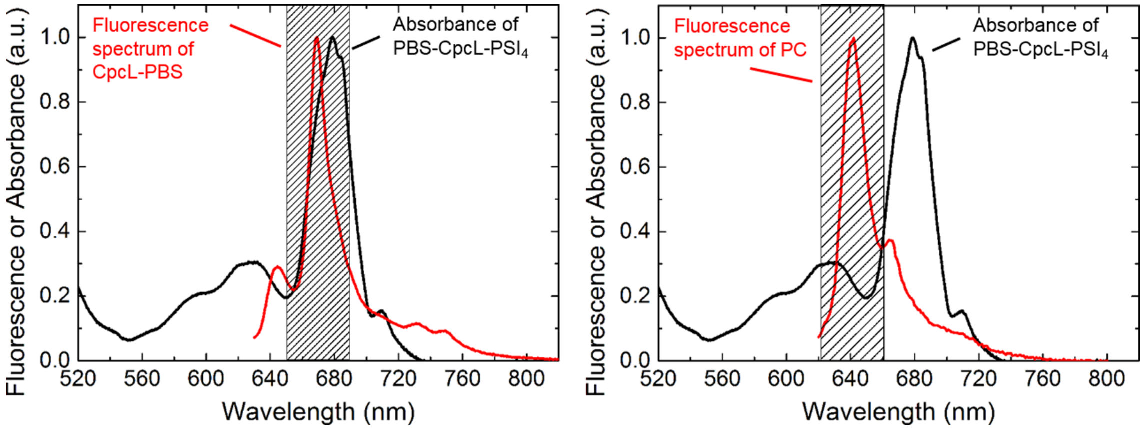


**Figure S9.** The overlap integral of fluorescence and absorbance at 77K. Red lines represent the emission spectra of CpcL-PBS (A) or PC (B). Black lines correspond to the absorption spectra of $PSI_4$. Fluorescence spectrum of PC in Ref. 1 was used. The $J$ value was calculated in a range from an emission peak to ± 20 nm (highlighted areas). The $J$ values of (A) and (B) are $7.1 \times 10^{13}$ $cm^6$ and $2.3 \times 10^{13}$ $cm^6$, respectively.

REFERENCES

(1) Watanabe, M.; Semchonok, D. A.; Webber-Birungi, M. T.; Ehira, S.; Kondo, K.; Narikawa, R.; Ohmori, M.; Boekema, E. J.; Ikeuchi, M. Attachment of Phycobilisomes in an Antenna-photosystem I Supercomplex of Cyanobacteria. *Proc. Natl. Acad. Sci. U. S. A.* **2014**, *111*, 2512-2517.

(2) Theiss, C.; Schmitt, F. J.; Pieper, J.; Nganou, C.; Grehn, M.; Vitali, M.; Olliges, R.; Eichler, H. J., Eckert, H. J. Excitation Energy Transfer in Intact Cells and in the Phycobiliprotein Antennae of the Chlorophyll *d* Containing Cyanobacterium *Acaryochloris marina*. *J. Plant. Physiol.* **2011**, *168*, 1473-1487.

(3) Pizarro, S. A.; Sauer, K. Spectroscopic Study of the Light-harvesting Protein C-phycocyanin Associated with Colorless Linker Peptides. *Photochem. Photobiol.* **2001**, *73*, 556-563.

(4) Watanabe, M.; Ikeuchi, M. Phycobilisome: Architecture of a Light-harvesting Supercomplex. *Photosynth. Res.* **2013**, *116*, 265-276.

(5) Mullineaux, C. W. Excitation-energy Transfer from Phycobilisomes to Photosystem-I in a Cyanobacterium. *Biochim. Biophys. Acta* **1992**, *1100*, 285-292.

(6) Byrdin, M.; Jordan, P.; Krauss, N.; Fromme, P.; Stehlik, D.; Schlodder, E. Light Harvesting in Photosystem I: Modeling Based on the 2.5-Å Structure of Photosystem I from *Synechococcus elongatus*. *Biophys. J.* **2002**, *83,* 433-457.

(7) Dixon, J. M.; Taniguchi, M.; Lindsey, J. S. PhotochemCAD 2: A Refined Program with Accompanying Spectral Databases for Photochemical Calculations. *Photochem. Photobiol.* **2005**, *81*, 212-213.

(8) Demidov, A. A.; Borisov, A. Y. Computer-simulation of Energy Migration in the C-phycocyanin of the Blue-Green-Algae *Agmenellum Quadruplicatum*. *Biophys. J.* **1993**, *64*, 1375-1384.

(9) Porra, R. J.; Thompson, W. A.; Kriedemann, P. E. Determination of Accurate Extinction Coefficients and Simultaneous Equations for Assaying Chlorophylls *a* and *b* Extracted with Four Different Solvents: Verification of the Concentration of Chlorophyll Standards by Atomic Absorption Spectroscopy. *Biochim Biophys Acta* **1989**, *975*, 384-394.
